# Modulation of naturalistic maladaptive memories using behavioural and pharmacological reconsolidation-interfering strategies: A systematic review and meta-analysis of clinical and ‘sub-clinical’ studies

**DOI:** 10.1101/282293

**Authors:** Katie H Walsh, Ravi K Das, Michael E Saladin, Sunjeev K Kamboj

**Affiliations:** Clinical Psychopharmacology Unit, Research Department of Clinical, Educational and Health Psychology, University College London, Gower Street, London, WC1E 6BT, UK; Department of Health Sciences and Research, College of Health Professions, Medical University of South Carolina, Charleston, USA

## Abstract

Consolidated memories can undergo enduring modification through retrieval-dependent treatments that modulate reconsolidation. This has been suggested to represent a potentially transformative clinical strategy for weakening or overwriting the maladaptive memories that underlie substance use and anxiety/trauma-related disorders. However, the ability to modulate *naturalistic* maladaptive memories may be limited by ‘boundary conditions’ imposed on reconsolidation by the nature of these memories. As such, the true potential of ‘reconsolidation therapy’ is currently unknown. Here, we report a meta-analyses of behavioural and pharmacological studies examining retrieval-dependent modulation of reward and threat memories in (sub)clinical substance use and anxiety/trauma respectively.

Of 4936 publications assessed for eligibility, 7 studies of substance use, and 9 of anxiety (phobia) and trauma-related symptoms were included in the meta-analyses. Overall, the findings were in the predicted direction, with the majority of effect sizes favouring the ‘Retrieval + Treatment’ condition. However, the magnitude of effects depended upon the nature of the treatment type, with pharmacological interventions (relative to behavioural strategies) showing a clearer beneficial effect in studies of phobia/trauma and post-retrieval behavioural strategies, a (significantly) larger effect in substance use studies. However, high levels of heterogeneity and small sample sizes limit the strength of conclusions that can be drawn at this stage of inquiry. We hope this review will provide an impetus to address these issues in future research.

## Introduction

### Phobia, Traumatic Stress and Substance Use Disorders as Disorders of Memory

Threat-related (phobia and traumatic stress-), and substance use disorders (SUDs) can be conceptualised as disorders of maladaptive associative memory (Fanselow & Sterlace, 2014; Hyman, 2005; McCarthy, Baker, Minami, & Yeh, 2011). The processes underlying formation and maintenance of these maladaptive memories are thus highly relevant to the treatment of these disorders. The failure of existing therapies to attenuate the emotional/motivational influence of maladaptive memories is one reason why treated individuals are vulnerable to relapse, even after prolonged remission/abstinence (Shaham, Shalev, Lu, De Wit, & Stewart, 2003; Shalev, Grimm, & Shaham, 2002; Staiger & White, 1991).

Recent advances in neuroscience have set the stage for the development of a new generation of treatments that focus on reducing the symptom-maintaining influence of maladaptive memories and the attainment of lasting protection against relapse (Kamboj & Das, 2017). The current review focuses on a specific memory retrieval-dependent form of memory plasticityreconsolidationthat can potentially be manipulated to ameliorate anxiety/trauma and SUD symptoms by targeting the *naturalistic* maladaptive memories that underlie them.

### Memory Reconsolidation

Historically, consolidated memories in long-term storage were thought to be stable and resistant to modification (cf., McGaugh, 2000). However, over the past two decades numerous studies have convincingly demonstrated that under certain retrieval conditions, even apparently long-established, putatively cortically-distributed memories can enter a transient labile state during which they are susceptible to modification before being re-stored in long-term memory (e.g. Gräff et al., 2015; Robinson & Franklin, 2010; Suzuki et al., 2004). This process is commonly referred to as *reconsolidation* and consists of two temporally and pharmacologically dissociable stages: (a) retrieval-induced *reactivation* or *destabilisation* of a previously consolidated memory and (b) its *restabilisation* in an updated or strengthened form Although reactivation engenders a period of memory instability which is required for normative memory strengthening and updating, stored representations are also susceptible to pronounced disruption during reconsolidation using pharmacological agents and behavioural procedures.

### Weakening maladaptive memories: Disruption of restabilisation with pharmacological agents

The restabilisation phase of the reconsolidation cycle is protein synthesis-dependent. Drugs that interfere (upstream or directly) with protein synthesis can therefore disrupt it. The most potent of these drugs (e.g. anisomycin or cyclohexamide) interfere directly with cellular translational machinery and macromolecule biosynthesis. However, these drugs are toxic and not safe for human use. As such, an alternative approach has involved indirect inhibition of protein synthesis through, for example, upstream neurotransmitter blockade. While a number of studies have examined such indirect modulation via diverse drugs (e.g. glucocorticoid, glutamatergic and GABAergic compounds), there are relatively few human studies using these drug classes (*c.f*. Das et al., 2015a; Drexler, Merz, Hamacher-Dang, & Wolf, 2016; Meir Drexler, Merz, Hamacher-Dang, Tegenthoff, & Wolf, 2015; Rodríguez et al., 2013; Wood et al., 2015). By contrast, the β-blocker, propranolol, has proven to be a particularly popular tool for probing reconsolidation in humans, especially in laboratory studies of fear conditioning (e.g. Bos, Beckers, & Kindt, 2014; Kindt, Soeter, & Vervliet, 2009; Schroyens, Beckers, & Kindt, 2017; Sevenster et al., 2013; Sevenster, Beckers, & Kindt, 2012a, 2014, Soeter & Kindt, 2015b, 2010, 2011, 2012a, 2012b). Other studies have extended these experimental findings with propranolol to clinical populations, showing enduring retrieval-dependent reductions in fear among people with specific phobia (Soeter & Kindt, 2015a), as well as drug craving among addicted individuals (e.g. Xue et al., 2017).

### Rewriting maladaptive memories using behavioural techniques

An alternative approach involves disrupting memory expression via reconsolidation interference using purely behavioural strategies (e.g. Monfils, Cowansage, Klann, & LeDoux, 2009). By targeting memory networks that are causally implicated in symptom expression, this approach aims to overcome the limitations of traditional inhibitory training (extinction) strategies. In particular, initially successful extinction is often followed by the ‘return of fear’or in the case of substance use disorders, the recurrence of craving and drug seeking-following re-exposure to unconditioned stimuli (USs; reinstatement), the simple passage of time (spontaneous recovery), or change in context (renewal). This strongly suggests that maladaptive associative memories persist following typical extinction-based therapies and might contribute to relapse (Bouton, 2002; Conklin & Tiffany, 2002). Reconsolidation-based behavioural (and pharmacological) treatments can potentially overcome these issues through a direct *updating* of reactivated memory networks.

In support of this, extinction learning after fear memory retrieval (so called ‘retrieval-extinction’) eliminates, and prevents the return of fear in rats (e.g. Monfils et al., 2009) and humans (Johnson & Casey, 2015; Schiller et al., 2009). Similarly, relative to extinction without prior retrieval, retrieval-extinction leads to enduring reductions in reactivity to drug cues in rats (e.g. Cofresí et al., 2017; Xue et al., 2012) and humans (Germeroth et al., 2017; Xue et al., 2012; see Kredlow, Unger, & Otto (2016) for a review of post-retrieval extinction effects), suggesting it is a general-purpose strategy for enduring modification of maladaptive memories. Other therapeutically applicable post-retrieval learning strategies might also be suited to updating appetitive and threat memories in humans, although these have received less attention (*cf* Das, Lawn, & Kamboj, 2015b; Hon, Das, & Kamboj, 2015).

### Putative boundary conditions on memory destabilisation

Despite the therapeutic implications of reconsolidation interference hinted at above, there appear to be some inbuilt limits on the regular destabilisation-restabilisation of naturally acquired memories. In particular, to minimise ongoing and indiscriminate memory interference, destabilisation following retrieval is constrained by a number of proposed ‘boundary conditions.’ Of particular relevance to naturalistic maladaptive memories, older and more strongly-encoded associations appear to be relatively resistant to destabilization following simple retrieval procedures (e.g. Alfei, Ferrer Monti, Molina, Bueno, & Urcelay, 2015; Milekic & Alberini, 2002; Robinson & Franklin, 2010; Suzuki et al., 2004). In contrast, experimental studies showing robust reconsolidation effects, particularly in humans, often involve experimentally-generated memories (especially conditioned fear), which are often reactivated mere days after training. These simulated maladaptive memories reflect profoundly different learning intensities compared to the naturalistic maladaptive memories found in phobia/trauma and SUDs. Associative learning in these disorders involves highly salient USs at encoding (supporting single trial learning) or reinforcement over many years in multiple contexts. For example, the typical ‘pack-a-day’ smoker, will experience close to 10^6^ reinforcements (puffs on a cigarette) over 12 years of regular smoking. These distinct properties of naturalistic memories (asymptotic learning and temporal remoteness) relative to experimentally-learned associations (sub-maximal learning and recency) potentially severely limit the application of findings from experimental conditioning studies to the treatment of some psychological disorders.

In addition, variation in stimulus predictability at retrieval may moderate the ability of retrieval procedures to labilise naturalistic maladaptive memories. In particular, accumulating experimental evidence suggests that a relevant prediction error (PE) at retrieval may be important for enabling full destabilisation of memory networks (e.g. Alfei et al., 2015; Exton-mcguinness, Lee, & Reichelt, 2015; Pedreira, Pérez-Cuesta, & Maldonado, 2004; Sevenster et al., 2013, 2014). As an illustration, Das and colleagues (2015b) found that while simple retrieval cues (followed by counterconditioning) produced intermediate levels of memory updating, incorporation of a PE at retrieval appeared to result in more pronounced rewriting of alcohol memories. As such, studies demonstrating weakening/updating of naturalistic maladaptive memories without the use of explicit PE-generating procedures during retrieval (the majority of studies) may reflect a lower bound of efficacy of such interventions, due to sub-optimal reactivation of maladaptive memory networks. However, while evidence of the PE-dependence of destabilisation has been demonstrated in fear conditioning experiments in humans (Sevenster et al., 2013), this has yet to be tested through systematic variations in degree of PE during reactivation of naturalistic memories associated with fixed and unknown learning histories. More generally, optimal retrieval parameters (e.g. duration or number of CS presentations at retrieval or the use of USs rather than CSs at retrieval; Exton-mcguinness et al., 2015) have not been thoroughly studied in humans, leaving some uncertainty about the suitability of the retrieval procedures used in extant studies of naturally acquired memories.

### The current review

To date, reviews and meta-analyses on reward and fear memory reconsolidation have either largely focused on non-human animals (e.g. Das, Freeman, & Kamboj, 2013) or, in the case of human studies, primarily on experimentally-generated memories, examining a single reconsolidation interference strategy (e.g. Kredlow et al., 2016; Lonergan, Brunet, Olivera-Figueroa, & Pitman, 2013) or memory system (Scully, Napper, & Hupbach, 2017). Such analyses are critical for furthering our understanding of the modulators of this fundamental memory process. However, a determination of the utility of reconsolidation modulation as a clinical strategy requires a synthesis of studies in which clinically-important symptoms are targeted in appropriate populations. To our knowledge, no comprehensive synthesis has been conducted on the effects of reconsolidation modulation strategies specifically directed at clinically relevant reward and threat-related memories in humans. The distinct properties of strongly encoded and remote naturalistic maladaptive memories versus those formed during experimental procedures may be extremely important in determining the translational utility of laboratory findings. Moreover, it might be that differences in the neural substrates of learning and distinctive learning histories associated with appetitive memories *versus* threat-related memories, render addictive and phobia/traumatic-stress disorders differentially susceptible to memory-modifying treatments due to differences in ‘reactivation-potential’ of their underlying maladaptive memories. However, this has yet to be formally tested. Finally, a systematic comparison of behavioural *versus* pharmacological strategies has not be conducted. The current meta-analysis addresses the lack of a systematic synthesis of behavioural pharmacological reconsolidation-interference strategies applied to human substance using and anxious/trauma-exposed (clinical and sub-clinical) samples.

## Methods

### Search Strategy

Psychinfo, PubMed, Web of Science, and Scopus databases were first searched on the 01/03/2017 and a renewed search conducted on 03/10/2017 using search terms based on a scoping search on experimental and therapeutic modulation of reconsolidation. The search terms were: (memory) AND ((((((((reactivat*) OR destabiliz*) OR destablis*) OR memory reconsolidation) OR reconsolidation) OR reconsolidation-extinction) OR extinction) OR retrieval) AND ((((((((((((((((((pharmacologic*) OR NMDA) OR N-methyl-D-aspartate) OR adrenoceptor) OR adrenergic) OR noradrenergic) OR beta adreno) OR adrenoreceptor) OR sympathetic) OR sympathetic nervous system) OR dopamine) OR dopaminergic) OR glucocorticoid*) OR cortisol) OR benzodiazepine) OR calcium channel) OR extinction) OR exposure) AND (((((((((((avers*) OR appetit*) OR fear) OR anxiety) OR PTSD) OR addiction) OR substance use disorder) OR substance use) OR drug use) OR drug) OR reward).

The search was limited to human studies and excluded reviews. The international clinical trials registry platform and *clinicaltrials.gov* were searched using the term “reconsolidation,” after which a search of the identified authors’ current publications was conducted. The reference lists of the following reviews were also checked for relevant studies: (Centonze, Siracusano, Calabresi, & Bernardi, 2005; de Kleine et al., 2013; de Quervain, Roozendaal, Nitsch, McGaugh, & Hock, 2000; Dennis, Perrotti, & Drug, 2015; Farach et al., 2012; Gisquet-Verrier & Riccio, 2012; Högberg, Hogberg, Nardo, Hallstrom, & Pagani, 2011; Kredlow et al., 2016; Lee, Nader, & Schiller, 2017; Makkar, Zhang, & Cranney, 2010; A. Milton, 2012; A. L. Milton & Everitt, 2010; Pitman, 2011; Schwabe, Nader, & Pruessner, 2014). Authors of all included studies were contacted regarding unpublished data.

### Study inclusion criteria

Figure 1 outlines the search, screening and section process, in line with the Preferred Reporting Items for Systematic Reviews and Meta-Analyses (PRISMA). Selection of studies was restricted to those that examined a reconsolidation-modulating (retrieval-dependent) pharmacological or behavioural strategy targeting naturally, rather than experimentally, acquired memories. In addition, studies were required to assess symptoms relevant to substance use or anxiety disorders reflecting effects on long-term (≥24 hr) memory. Participants were required to be recruited on the basis of elevated anxiety, experience of trauma or problematic alcohol/substance use. There was no requirement for a formal diagnosis or for participants to be seeking treatment. Studies were required to randomise adult participants to a Retrieval + (reconsolidation interfering) ‘Treatment’ or control group (see below) and contain n≥15 per condition at randomisation. Only studies reported in English were included. Abstracts were reviewed for eligibility by the first author. Sixteen studies that examined pharmacological or behavioural strategies for modifying naturalistic appetitive or threat-related memories via reconsolidation in clinical or subclinical human samples were included. Note, one study (Jobes et al., 2015) that initially met inclusion criteria was excluded following discussion due to the complex nature of the design, which involved participants receiving methadone at various times during the intervention (either pre- or post-reactivation). This was in addition to the specific reconsolidation-interfering study-medication (propranolol), making it impossible to disentangle opioid from β-adrenergic treatment effects. In addition, it should be noted that a recent study on the effects of propranolol on smoking memories (Xue et al., 2017) did not meet criteria because the effects primarily related to experimentally acquired, rather than naturalistic smoking memories.

**Figure 1.**
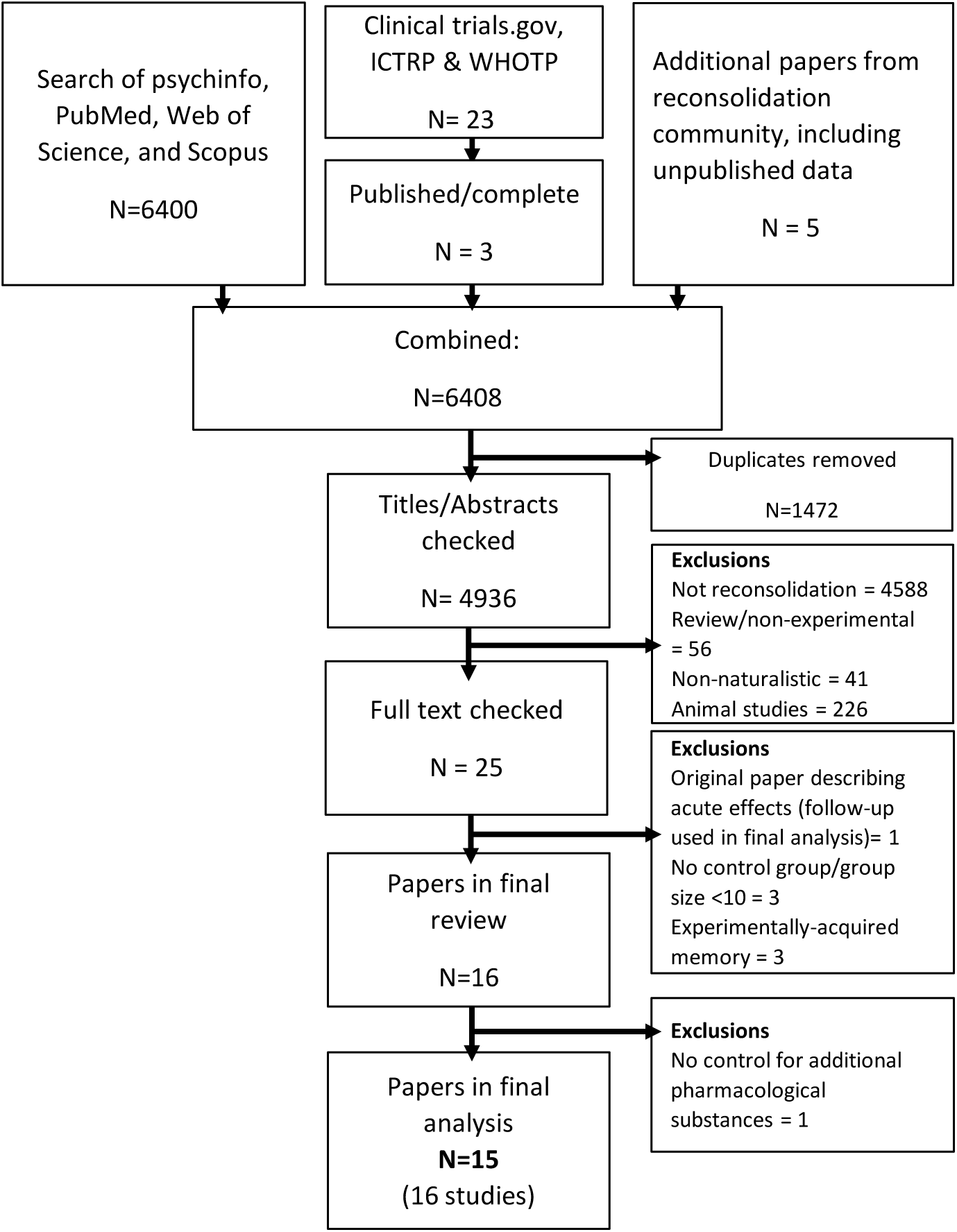
PRISMA flowchart of study inclusion process

### Methodological evaluation of studies

Identifying information (authors, institutions, journal details and significance of results) was removed from included papers, allowing anonymised methods sections to be independently assessed by two nominally blind investigators. The tool for methodological appraisal was a modified version of an instrument used in our previous meta-analysis of reconsolidation studies (Das et al., 2013). The level of inter-coder agreement was 83% and any discrepancies in ratings were resolved through discussion.

### Data Extraction

Details regarding the study protocol, including memory retrieval procedure, outcome measures, treatment timing (relative to retrieval), type (behavioural or pharmacological) and dose, as well as ‘disorder’ type were extracted from the selected articles.

### Outcome measures

A preliminary review of the selected studies identified specific outcomes for use in effect size (ES) calculation. These were selected based on the regularity with which these measures were reported across studies. We chose this approach in preference to determining ESs for published significant effects in order to minimise bias, since some of the included studies were not identified as clinical trials, and therefore had no pre-determined (registered) outcomes. As such, subjective craving – an important clinical target in SUD treatment, reflecting conditioned responding to drug cues (i.e. the subjective expression of retrieved drug-related memories) was the primary outcome in the current analysis of substance use studies, as it was reported in all relevant publications. Similarly, studies of phobias consistently used the behavioural approach (avoidance) test (BAT), although the nature of outcomes from this test varied from study to study (e.g. distance between participant and feared object, Shiban, Brütting, Pauli, & Mühlberger, 2015; subjective fear ratings during proximal approach, Telch, York, Lancaster, & Monfils, 2017). Finally, trauma-related studies most commonly reported PTSD symptom severity (three studies; *Table 1*), apart from one study, which reported memory performance (number of recalled trauma event details; Kredlow & Otto, 2015).

**Table 1.**
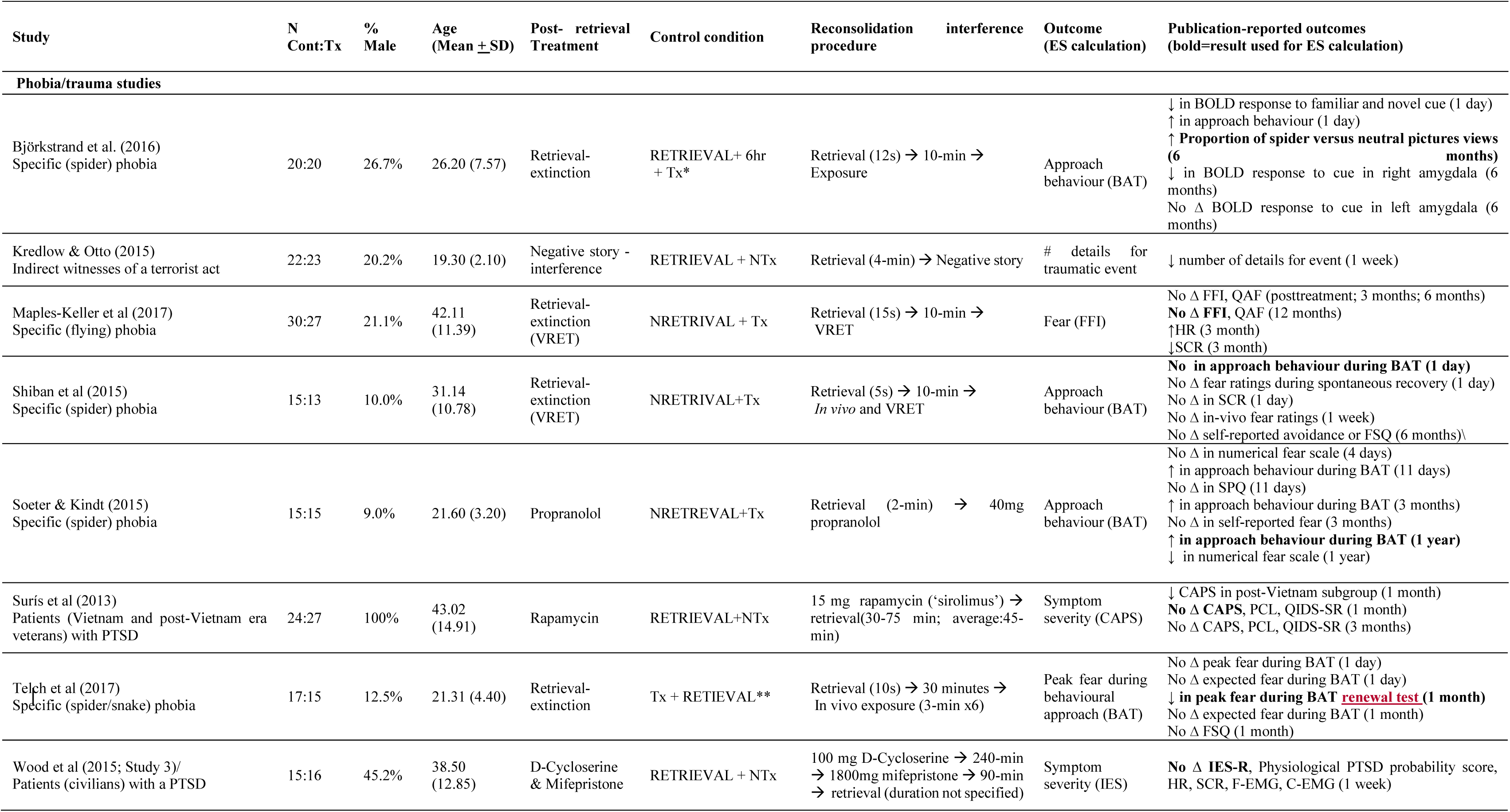
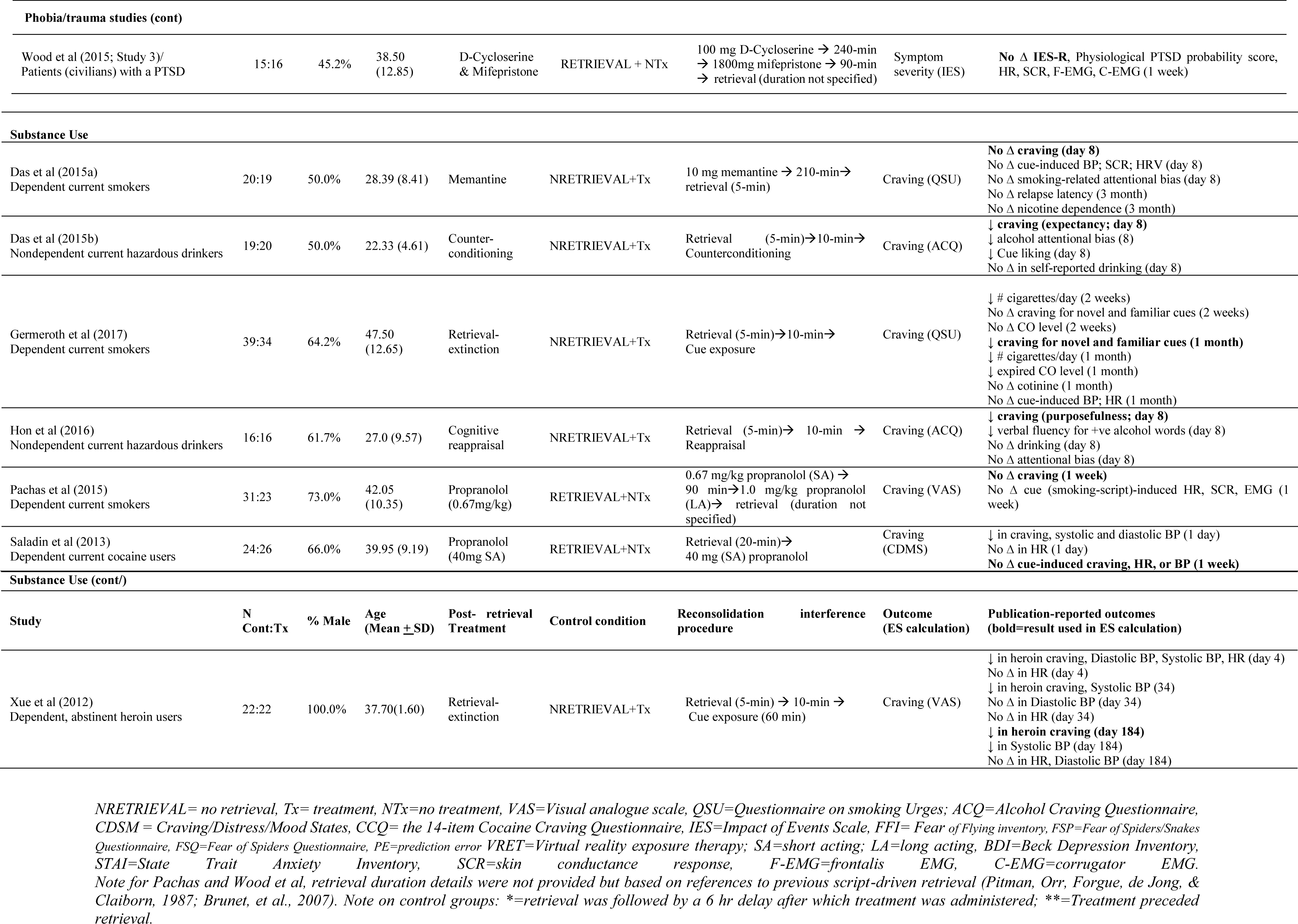
Characteristics of studies

### Statistical approach

#### Effect size determination

Data required for effect size (ES) determination were extracted and entered into an Excel spreadsheet by the first author. Random effects models (Dersimonian & Laird, 1986) were selected and the generic inverse variance method used. ESs were calculated as between groups standardised mean differences (Hedge’s g; Higgins & Green, 2011) using the Review Manager Software (version 5.3; The Cochrane Collaboration, 2014) and interpreted using the standards of Cohen (1988) and Sawilowsky (2009): ∼0.1=very small, ∼0.2=small; ∼0.5=medium; ∼0.8=large and ∼1.2=very large. Intermediate descriptive labels (e.g. small-medium) were used to describe ESs, where appropriate.

ESs related to the primary 1 *df* comparison of interest, namely, Retrieval + Treatment (pharmacological or behavioural) versus a suitable control condition. A comparison with a No Retrieval + Treatment control was deemed to best represent the specific effect of a memory interfering/weakening treatment via reconsolidation. Where such a group was not used, ESs were calculated relative to a Retrieval + No Treatment condition. Other control groups are also suitable for testing reconsolidation effects. Unlike pharmacological studies, in which drug effects are likely to be present for several hours (i.e. during the period of memory lability) even if the treatment is administered prior to reactivation, retrieval-dependent memory-interfering behavioural treatment effects are theoretically constrained if the treatment occurs before retrieval. As such, Treatment *followed* Retrieval is a suitable control condition in behavioural studies (however see Hutton-Bedbrook & McNally, 2013) for discussion of effects that are not consistent with a standard reconsolidation interpretation). Finally, treatment delivered outside of the ‘reconsolidation window’ were selected as controls for memory reactivation as this is a time limited process, such that treatment delivered outside of the window will not modify the original memory, which returns to an inactive state in the hours after reactivation (i.e. <6 hr after retrieval).

Given that reward- and threat-related disorders have distinct multipath aetiologies and underlying learning processes, these disorder types were evaluated separately in meta-analyses. Alternatively, given the aetiological similarity in terms of the proposed central role of classical conditioning in specific phobias and trauma-related disorders, these two classes of disorders were considered together as a single category (phobia/trauma). Further, using subgroup analysis, we examined whether treatment type (behavioural versus pharmacological) produced different population ES estimates within each broad disorder type. Finally, we examined moderation by gender ratio, participant age and score on the methodological appraisal tool (based on number of positively endorsed desirable study characteristics as a proportion of the total number of items that could be positively endorsed) across all *studies* using these as continuous variables in meta-regressions. Note, although variation in retrieval parameters (especially retrieval trial duration and time between reactivation and treatment) could affect the extent to which memories are reactivated or weakened/over-written, insufficient variability in, and lack of these details about these parameters prevented us from exploring these as moderators (*cf* Kredlow et al., 2016).

Subgroup analyses and forest plots were derived from RevMan. Heterogeneity across studies was assessed using the *I*^*2*^ statistic and described qualitatively thus: ∼25% =low; ∼50%=moderate, ∼75% = high (Higgins, Thompson, Deeks, & Altman, 2003). Sensitivity analysis was conducted when heterogeneity was high and involved sequentially testing the effects of (removing) individual studies to determine which had the greatest influential on heterogeneity. Alternative aggregate ESs are reported where removal of the most influential study resulted in a reduction of heterogeneity to moderate levels or below (i.e. *I*^*2*^*<50%*).

Where insufficient information was available in publications to calculate ESs from means/SDs and these details were not available from authors (Pachas, Gilman, Orr, Hoeppner, Carlini, Loebl, et al., 2015; Xue et al., 2012) estimates were obtained from figures in the relevant publications using Plot Digitizer software (Poisot, 2011). Publication bias (symmetry of funnel plots and trim and fill) was assessed using the MAVIS package Version 1.1.3 (Hamilton, Aydin, & Mizumoto, 2017).

#### Terminology

We use the terms ‘destabilisation’ and ‘reactivation’ interchangeably to refer to a behaviourally silent memory state whose occurrence can only be inferred indirectly through observed memory modification. ‘Retrieval’ is used here to refer to experimental procedures that are intended to reactivate/destabilise memory, but which may or may not be successful in this regard. The term ‘retrieval’ is not intended to imply recall of a discrete memory trace (*cf* Telch et al., 2017), but rather, retrieval or reactivation of a more complete *network* of reward (substance use) or aversive (phobic/trauma-related) associations.

## Results

### Study and sample characteristics

After exclusions, the literature search yielded a total of 16 studies from 15 publications (n=673). Five were studies on specific phobias, four on trauma-related symptoms and seven studies examined substance use. Of these, four examined post-retrieval pharmacological interventions (Soeter & Kindt, 2015a; Surís, Smith, Powell, & North, 2013; Wood et al., 2015) studies 2 and 3) and five, behavioural strategies (Björkstrand et al., 2017; Kredlow et al., 2016; Maples-keller et al., 2017; Shiban et al., 2015; Telch et al., 2017). The seven substance use studies also examined either pharmacological (*k*=3; Das et al., 2015a; Pachas et al., 2015; Saladin et al., 2013) or behavioural reconsolidation interference strategies (*k*=4; Das et al., 2015b; Germeroth et al., 2016; Hon et al., 2015; Xue et al., 2012).

Participant details (gender ratio; age) are presented in *Table 1*. There was considerable variation between studies in terms of gender ratio of participants. Among substance use studies, gender was generally balanced or had a higher proportion of men, in line epidemiological studies (Seedat et al., 2009). Xue et al (2012) was an exception as it only included detoxified male heroin users. In contrast, studies of phobia/trauma studies were generally skewed towards a higher representation of women, again, in line with epidemiological evidence (McLean, Asnaani, Litz, & Hofmann, 2011). An exception was the study by Surís et al (2013), which only recruited men (combat veterans). Participant age varied widely across studies, although the mean age of participants was not statistically different (p=0.57) in phobia /trauma studies (*M* =32.1, *SD* =10.5) and substance use studies (*M* =37.7, *SD* =9.2).

### General study methodologies

Key design features of studies and the presence/absence of specific desirable methodological study features are outlined in Tables 1 and 2. Table 2 shows that studies generally contained many desirable methodological features. The most common methodological limitations across studies were a lack of comprehensive experimental conditions that controlled for the effects of simple retrieval or treatment alone. In addition, a lack of experimenter/assessor blinding was a virtually universal limitation of the behavioural studies, but uncommon in pharmacological studies.

**Table 2.** Methodological/reporting features of studies.

| Study Name | a | b | c | d | e | f | g | h | i | j | k | l | M |
| --- | --- | --- | --- | --- | --- | --- | --- | --- | --- | --- | --- | --- | --- |
| Das., et al., 2015a | Y | Y | N | Y | Y | N | Y | Y | N/A | Y | Y | Y | Y |
| Das et al., 2015b | Y | Y | Y | N | Y | N | N | Y | N/A | Y | Y | N/A | Y |
| Germeroth, et al., 2017 | Y | Y | Y | N | Y | Y | N | Y | N/A | Y | Y | N/A | N |
| Hon et al., 2016 | Y | Y | N | N | Y | Y | N | Y | Y | N | N | N/A | Y |
| Pachas, et al., 2015 | Y | Y | N | Y | Y | Y | N | Y | N/A | N | Y | Y | N |
| Saladin, et al., 2013 | Y | Y | Y | Y | Y | Y | N | Y | N/A | Y | Y | Y | N |
| Xue, et al., 2012 | Y | Y | N | N | Y | Y | N | Y | N/A | Y | Y | N/A | Y |
| Soeter & Kindt, 2015 | Y | Y | N | N | Y | Y | Y | Y | N | Y | Y | Y | N |
| Telch et al., 2017 | Y | Y | Y | N | Y | Y | Y | Y | N/A | Y | Y | N/A | Y |
| Björkstrand et al., 2016 | Y | N | N | N | Y | N | N | N | N/A | Y | Y | N/A | Y |
| Shiban et al., 2015 | Y | Y | Y | Y | Y | N | N | Y | N/A | Y | Y | N/A | N |
| Maples-Keller et al., 2017 | Y | Y | N | Y | Y | Y | N | Y | N | Y | Y | N/A | Y |
| Kredlow & Otto, 2015 | Y | N | Y | Y | Y | Y | N | Y | Y | Y | Y | N/A | N |
| Surís et al., 2013 | Y | Y | N | Y | Y | Y | N | Y | N | Y | Y | N | Y |
| Wood, et al., 2015 (Study 2) | Y | Y | N | Y | Y | Y | Y | N/A | Y | N | Y | Y | Y |
| Wood, et al., 2015 (Study 3) | Y | Y | N | Y | Y | Y | N | Y | Y | N | Y | Y | Y |

*Y=Desirable study characteristic present; N=desirable characteristic not present; N/A=not applicable.*
A) Is the study design (or paradigm) described? B) Are there clear inclusion and exclusion criteria? C) Are the procedures for randomisation described? D) Are the procedures for blinding (If appropriate, i.e. if outcome is experimenter rated) described? E) Is/are primary outcome measure(s) clearly specified? F) Are participant demographics (i.e. age/gender) provided? G) Is there sufficient control? i.e. both a No Retrieval + Treatment and a Retrieval + No Treatment group. H) Were groups comparable at baseline? I) Where outcome measure was experimenter rated, was inter-rater reliability achieved and evaluated? J) Is length of retrieval trial given? K) Is treatment 'dose' given? L) Is timing of drug administration relative to reconsolidation clearly described? M) Was there a limited amount of missing data (<20%) or, if there was significant missing data, was this dealt with appropriately?

### Retrieval procedures

Most studies used *in vivo* exposure to CSs (e.g. powder resembling crack cocaine; a live spider), other visual representations of the CS (e.g. video of cocaine use; a series of pictures of spiders), or both, to reactivate memories. All of the trauma-related studies encouraged autobiographical recall of the traumatic incident(s) to reactivate trauma memory. Other studies also incorporated instructions to recall specific relevant autobiographical episodes evoked by the CSs (e.g. Das et al., 2015a, 2015b; Hon et al., 2015; Pachas et al., 2015; Telch et al., 2017) and four studies explicitly included prediction error at retrieval (Das et al., 2015a, 2015b; Hon et al., 2015; Soeter & Kindt, 2015a). The latter involved some form of expectation violation (e.g. generating an expectation that the participant will experience the US, and then violating this expectation; (Das, Gale, Hennessy, & Kamboj, 2018).

As outlined in Table 1, most studies specified the duration of the retrieval procedures used. The modal duration in substance use studies was 5 min (used in five of the six studies specifying retrieval duration); one study used a longer retrieval procedure (2 x 10 min; Saladin et al., 2013). Seven of the nine phobia/trauma studies specified the duration of the retrieval procedures, which varied more than the substance use studies. All of the phobia studies used ≤2 min retrievals, with most studies clustered in the 5-15s range. The two trauma-related studies that specified retrieval duration used 4 min, and 45 min.

### Pharmacological and behavioural reconsolidation interference procedures

Pharmacological studies most commonly used propranolol (*k* =2 substance use studies: Pachas et al., 2015; Saladin et al., 2013; k=1 phobia/trauma study: Soeter & Kindt, 2015a. The reconsolidation interfering effects of mifepristone (*k* =2; Wood et al., 2015; study 2 and 3) and sirolimus (rapamycin; *k* =1; Surís et al., 2013) on threat memory and memantine on reward memory (*k* =1; Das et al., 2015a) were also examined. In all cases, selection of these drugs by study authors was based on their putative downstream protein synthesis inhibiting effects, and particularly their tendency to interfere with the protein synthesis-dependent restabilisation phase of reconsolidation.

Among behavioural studies, retrieval-extinction was the most commonly tested procedure, either using ‘standard’ in vivo and/or picture-stimulus exposure for specific phobia (Bjorkstrand et al., 2016; Telch et al., 2017) and substance use (Germeroth et al., 2017; Xue et al., 2012) or virtual reality exposure for specific phobia (Maples-keller et al., 2017; Shiban et al., 2015). The remaining behavioural studies examined post-retrieval counterconditioning (Das et al., 2015b) and cognitive reappraisal (Hon et al., 2015) in substance using (heavy alcohol drinkers), or prose interference (Kredlow & Otto, 2015) in sub-clinical, trauma-exposed individuals.

### Study Outcomes

Across all studies, 13 of the 16 ESs were positive (favouring retrieval-dependent reconsolidation-interference). In all cases, ESs were based on comparisons between the retrieval and control condition on the last assessed time-point for the relevant outcome. This ranged from one day (Shiban et al., 2015), or more commonly, one week (Das et al., 2015b; Hon et al., 2015; Kredlow & Otto, 2015; Saladin et al., 2013; Wood et al., 2015) to 12 months (Soeter & Kindt, 2015a). We deemed this relatively stringent longest time-point comparison to be appropriate given the claim for *permanent* memory modification following reconsolidation interference.

Among the phobia/trauma studies, other than outcomes from the BAT test (phobia studies) and trauma symptom severity/trauma memory recall (trauma-related studies) used to calculate ESs, some of the reviewed publications reported additional outcomes showing significant retrieval-dependent benefits (Table 1). These included reduced skin conductance in response to fear-provoking stimuli (Maples-Keller, et al., 2017), subjective fear/phobic symptoms (Soeter & Kindt, 2015a) and neural activity in the amygdala (Björkstrand et al., 2017). Notably, reduction in fear (of spiders) in (Soeter & Kindt, 2015a) study only emerged at long-term follow-up, suggesting a lagged benefit in the retrieval group. Conversely, Maples-keller et al. (2017) reported *higher* physiological arousal (heart rate) at 3 month follow up in the retrieval group (at baseline *and* during exposure to feared cues) relative to the no retrieval control. However, this was interpreted as a relative benefit to the retrieval group (i.e. high levels of fear were thought to attenuate physiological reactivity in the no retrieval group, although there were no differences in fear ratings between groups).

In addition to craving, other statistically significant effects were also reported in a number of the substance use studies (Table 1). These included reductions in smoking (Germeroth et al., 2017), alcohol attentional bias (Das et al., 2015b), alcohol cue liking (Das et al., 2015b), fluency for positively valenced alcohol words (Hon et al., 2015), and cocaine and heroin cue-evoked blood pressure changes (Saladin et al., 2013; Xue et al., 2012).

### Control conditions

A control group that received treatment in the absence of putative reactivation (No Retrieval + Treatment) was considered the most appropriate comparison condition and was the most commonly employed. A number of pharmacological studies used a Retrieval + no Treatment (placebo) group (Pachas et al., 2015; Saladin et al., 2013; Surís et al., 2013; Wood et al., 2015) Study 3), as did (Kredlow & Otto, 2015), who compared negatively valenced interfering prose with a no prose condition.

### Effect size for symptoms of phobia and trauma

The aggregate ES for phobia/trauma symptoms was moderate (*k* =9; *n*=342; *g*=0.47, 95% CI [0.12, 0.81], *p* =0.01; *Figure 2*) and showed moderate heterogeneity (*I*^2^ = 59%). It is clear from inspection of the forest plots however, that Soeter & Kindt’s (2015a) study contributes disproportionately to the overall ES. A sensitivity analysis showed that exclusion of Soeter and Kindt (2015a) essentially eliminated heterogeneity (*I*^2^=5%), but also reduced the ES (*g*=0.34, 95% CI [0.11, 0.57]), although it remained significantly greater than 0 (p=0.004).

**Figure 2.**
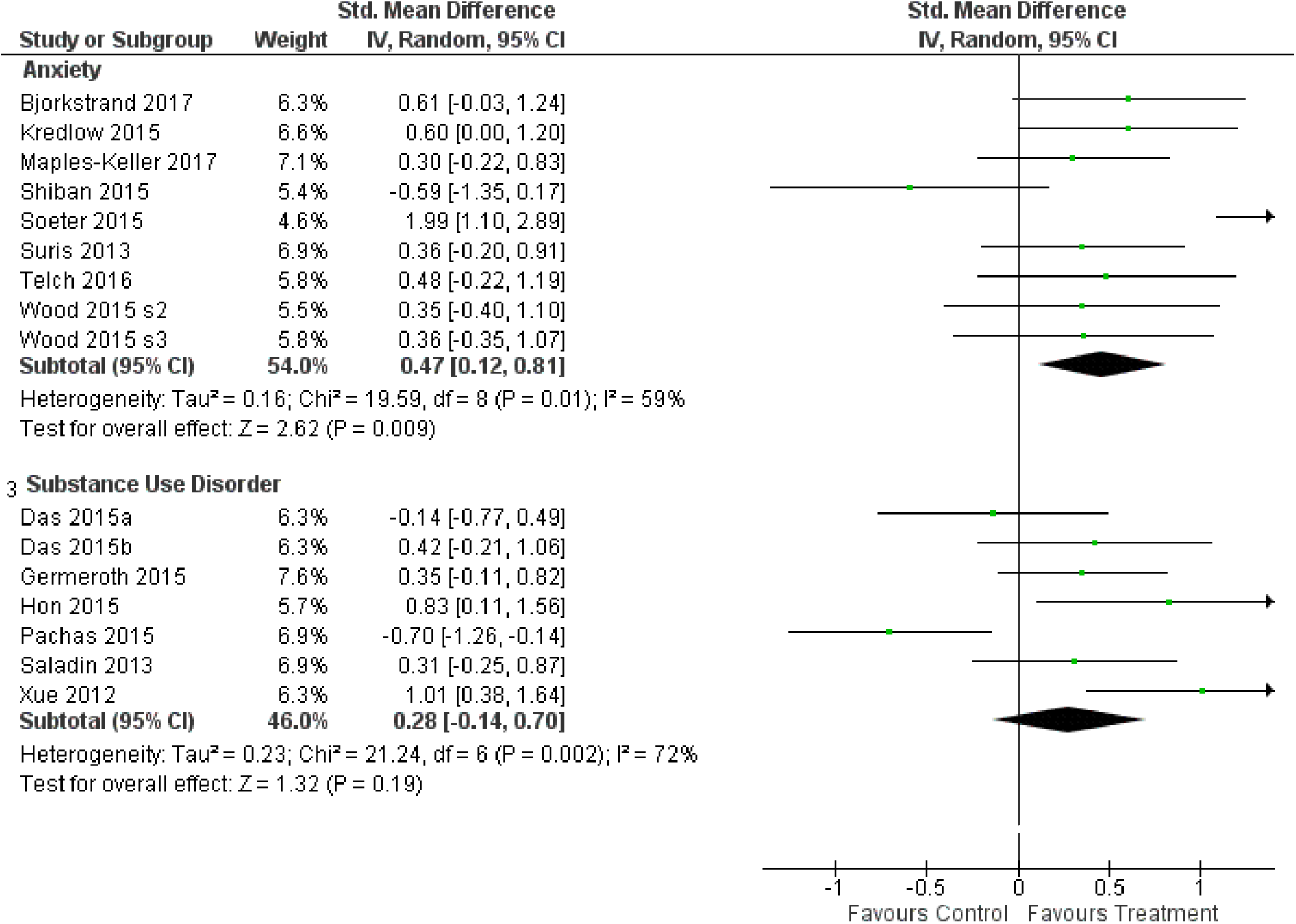
Forest plot of all included studies and a comparison of overall ES and associated CI and for anxiety and substance use studies.

The aggregate ES for all pharmacological studies was medium-large, although the confidence intervals for this estimate were wide (*k*=4, *g=*0.71, 95% CI [0.03, 1.39], p=0.04). Heterogeneity was correspondingly high (*I*^*2*^=72%). When Soeter and Kindt (2015a), was retained in the analysis, the population ES estimate had poor precision, with the true value lying in the range from no effect to a very large effect. Exclusion of Soeter & Kindt (2015a) completely eliminated heterogeneity (*I*^*2*^=0%) but also halved the ES, rendering it non-significant (*k=*3, *g* =0.36, 95% CI [-0.02, 0.74], *p*=0.06). The latter ES was similar to that of behavioural studies of phobia/trauma (*k=* 5, *g*=0.32, 95% CI [-0.06, 0.70]), which was also non-significant (p=0.13) and associated with a moderate degree of heterogeneity *(I*^*2*^*=43%)*. Subgroup analysis showed that pharmacological and behavioural studies did not significantly differ, regardless of the inclusion (χ^2^(1)=0.98, p=0.33) or exclusion (χ^2^(1)=0.02, p=0.90) of Soeter and Kindt (2015a).

### Effect size for symptoms related to substance use

Across all substance use studies, the aggregate ES was small and non-significant, with high levels of heterogeneity (*I*^*2*^ =72%). The confidence intervals covered a relatively wide range between a very small negative and a medium positive effect (*k* =7; *n*=331; *g=*0.28, 95% CI [-0.14, 0.70], *p* =0.19; *Figure 2*). Sensitivity analysis identified a single study (Pachas et al., 2015) that appeared to be especially influential. Its removal reduced heterogeneity to *I*^*2*^ =36% and increased the aggregate ES to a significant and small-moderate magnitude (*k*= 6; n=277; *g=* 0.44, 95% CI [0.14, 0.75], p=0.005).

Subgroup analysis of substance use studies indicated that pharmacological studies (including Pachas et al., 2015) were associated with a small, non-significant, negative ES (*k* =3; *n* =143; 0.18, 95% CI [-0.77, 0.42], *p*=0.56), with moderate-high levels of heterogeneity (*I*^2^ = 68%). The two pharmacological studies other than (Pachas et al., 2015) had a very small combined ES (*g* =0.11, 95% CI [-0.32, 0.55]). In contrast, behavioural studies yielded a significant, medium effect (*k*=4; *n* =188; 0.60, 95% CI [0.29, 0.92], *p* <0.001), with negligible heterogeneity (*I*^2^ = 11%). A moderator analysis (including Pachas et al., 2015) suggested that the ESs of behavioural and pharmacological studies of substance use were significantly different (χ^2^(1)=5.15, p=0.02; *Figure 3*).

**Figure 3.**
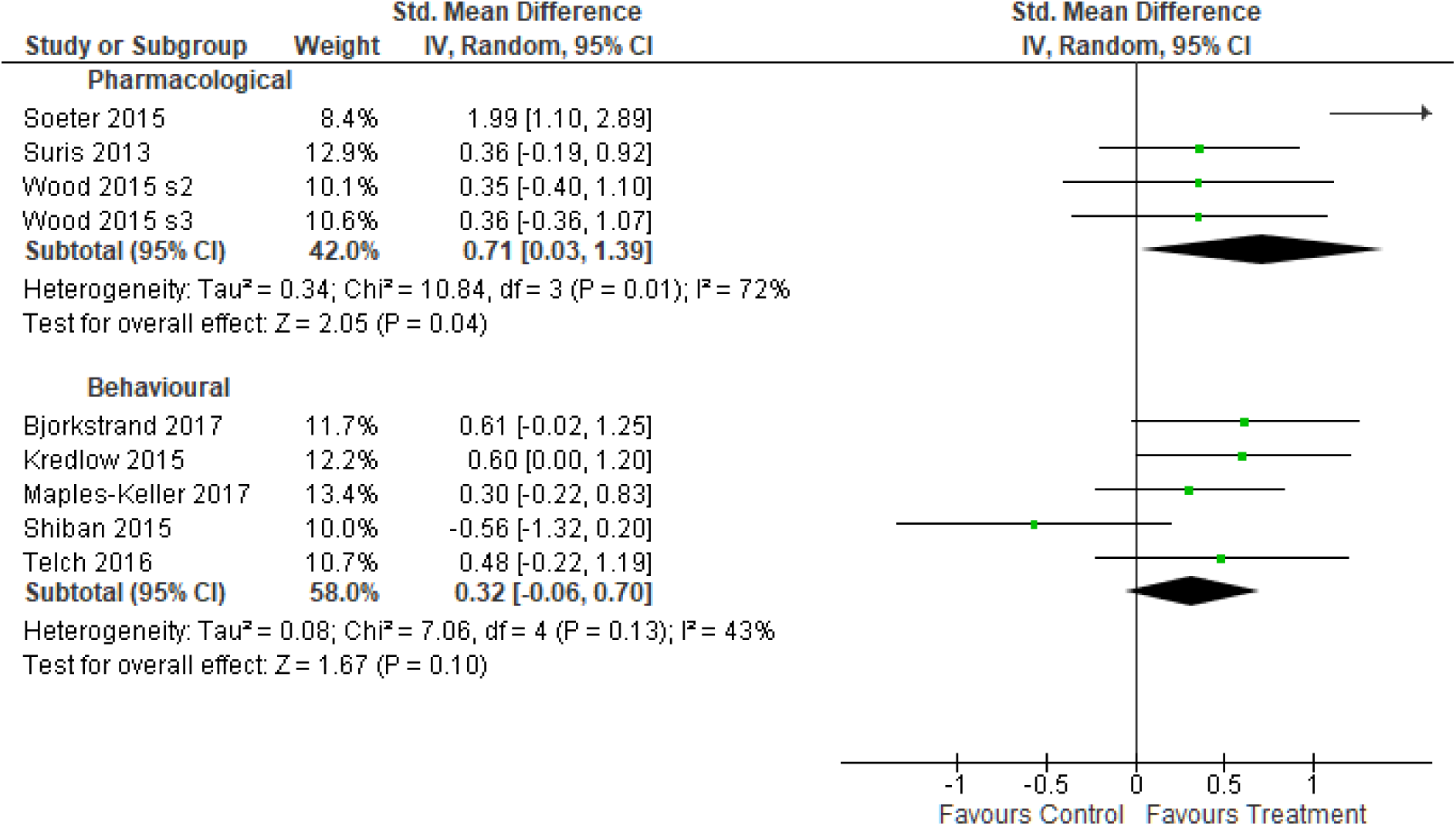
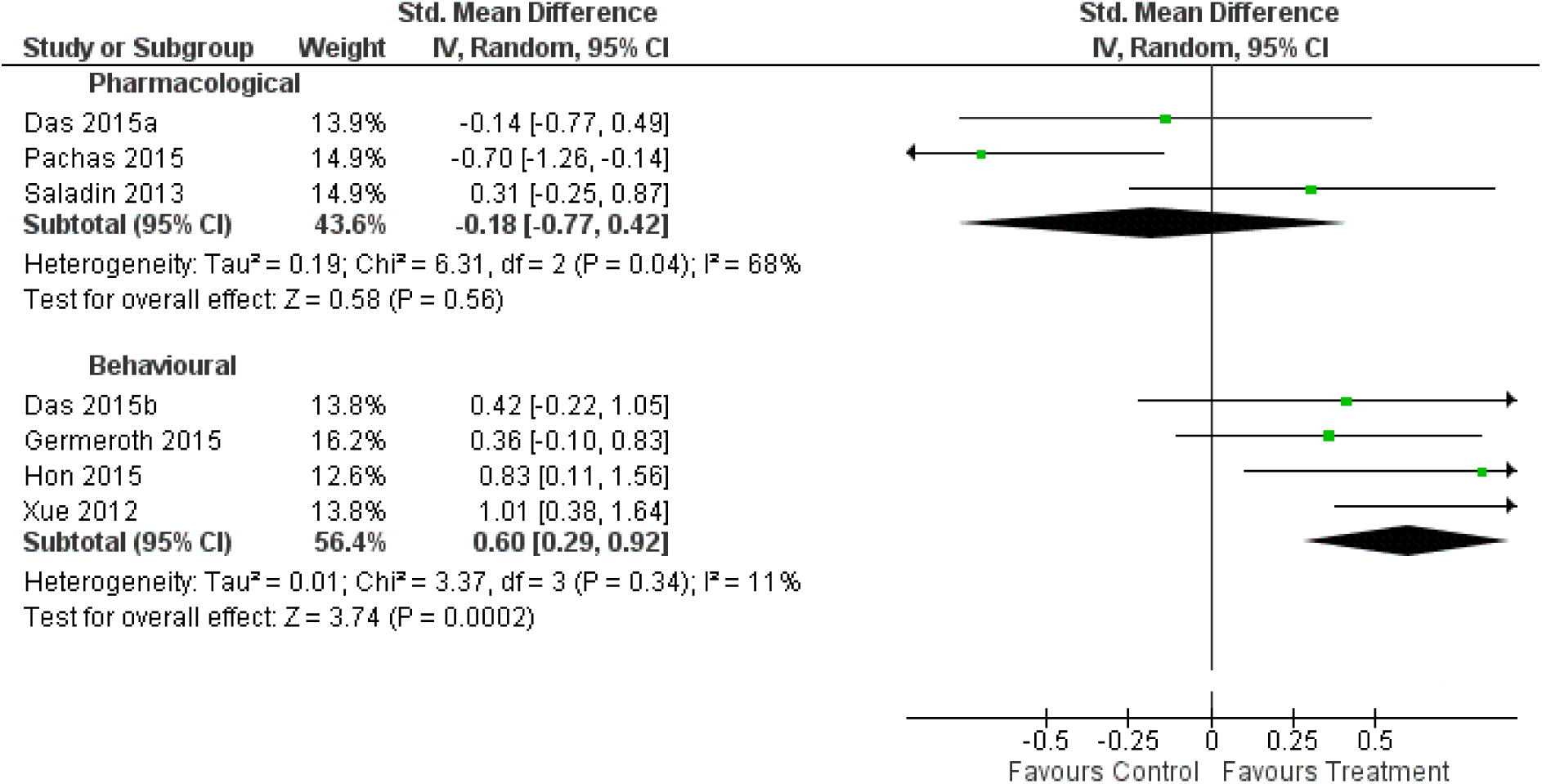
**a.** Comparative forest plot for treatment type (behavioural vs. pharmacological) in studies of maladaptive threat memories (anxiety) **b.** Comparative forest plot for treatment type (behavioural vs. pharmacological) in studies of reward memories (substance use)

### Moderation

Across *all* studies, meta-regression suggested that none of the specified moderators (age, proportion of female participants or methodological appraisal score) were significant predictors of ESs (*t* values <1.5, p values >0.1).

### Publication bias

A funnel plot for the phobia/trauma studies did not indicate asymmetry (*t* (7)=0.78, p=0.46; *Figure 4*). No adjustments to the effect of phobia/trauma studies was suggested by trim and fill (Duval & Tweedie, 2000a, 2000b). The funnel plot for studies of substance use similarly indicted a lack of asymmetry (t(6)= 0.3564, *p*=0.73; *Figure 5*), with no adjustments to effect following use of trim and fill. Overall, these results suggest an absence of publication bias for phobia/trauma and substance use studies.

**Figure 4.**
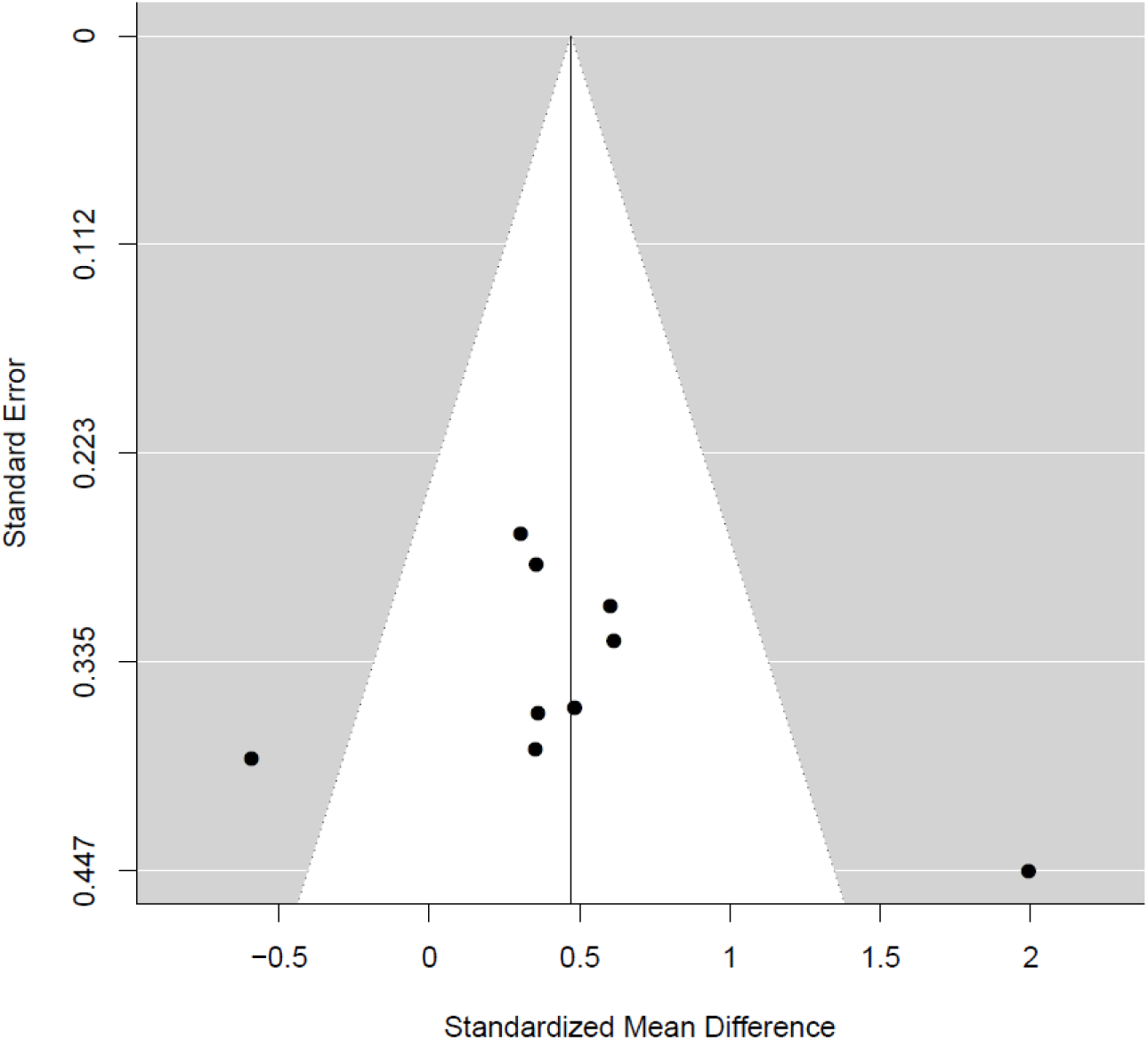
Funnel plot of ES against standard error for studies of anxiety/trauma.

**Figure 5.**
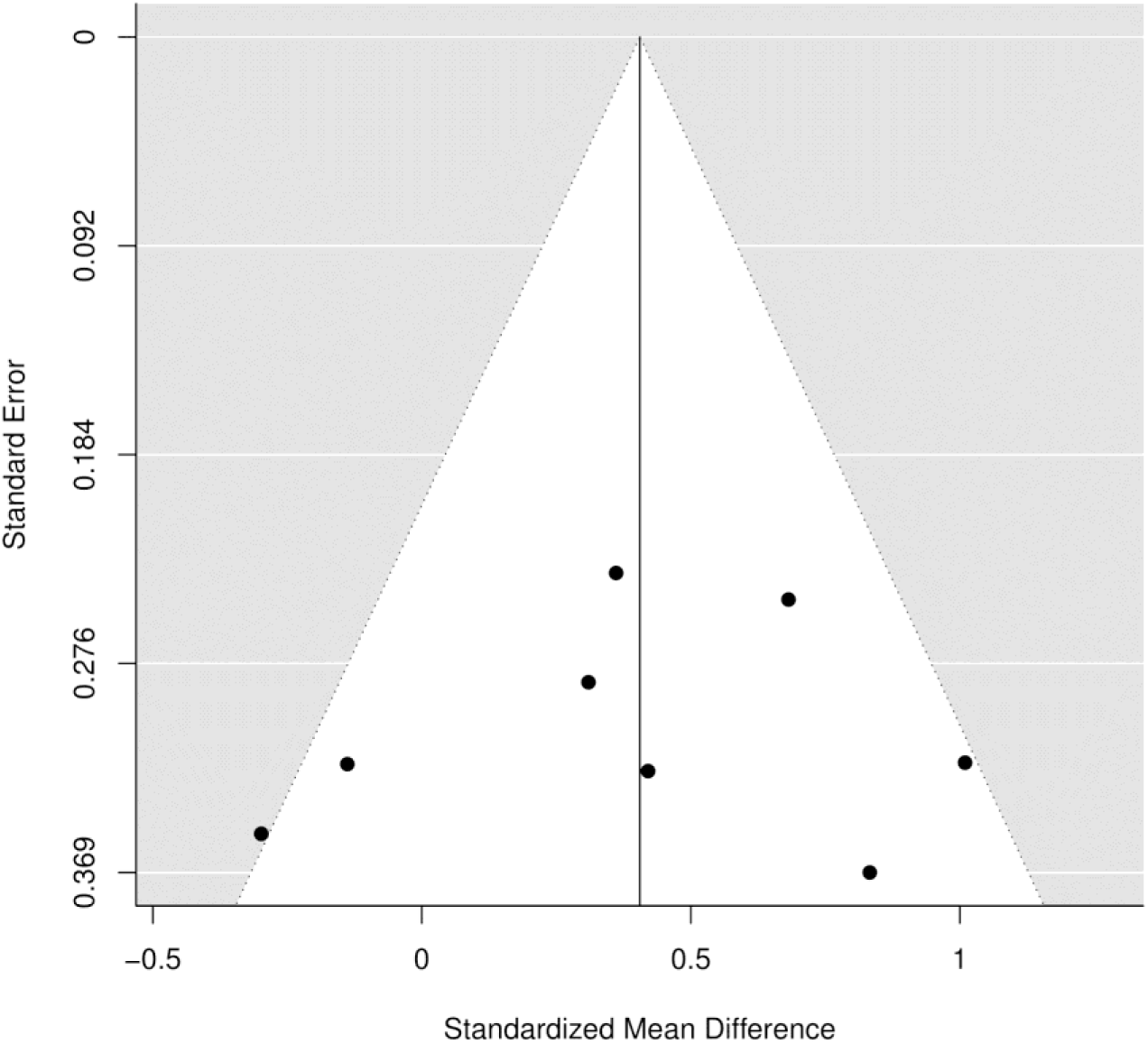
Funnel plot of ES against standard error for studies of substance use disorder.

## Discussion

This meta-analysis provides a synthesis and critical evaluation of research on reconsolidation of naturalistic maladaptive memories using pharmacological and behavioural memory-weakening/interference strategies in (sub)clinical samples. Extension of non-human and human experimental findings to clinically relevant populations is a relatively new area of translational research, with the oldest publication in this review dated 2012. As such, there are currently a small number of relevant studies, although findings across these were relatively consistent. In particular, 13 of 16 ESs were in the predicted direction (i.e. favouring a reconsolidation-modulation interpretation), but of the two broad disorder categories, only the population ES estimate for phobia /trauma (behavioural and pharmacological studies) was significant. Moderator analysis by intervention type (pharmacological versus behavioural) indicated larger effects in behavioural versus pharmacological studies in the case of substance use, while the opposite pattern was observed for phobia/trauma studies (although the comparison was not statistically significant in the latter case). However these general findings need to be considered in the context of population ES estimates for both phobia/trauma and substance use studies that were each substantially influenced by a single study which inflated heterogeneity, and skewed the results towards either a larger (phobia/trauma) or smaller (substance use) overall ESs.

### General overview of studies

Across all substance use studies, the overall ES was small and non-significant, although removal of (Pachas et al., 2015) reduced heterogeneity and increased the aggregate ES estimate, rendering it significant. Anxiety/trauma studies had a significant medium ES when all studies were considered, and a small, but still significant effect when Soeter and Kindt (2015a) was excluded. Overall, these findings support the idea that, despite their chronological age and strength relative to experimentally acquired memories, naturalistic maladaptive memories are capable of being destabilised and subsequently weakened/over-written using reconsolidation-modulating strategies. As such, the putative boundary conditions (memory remoteness and strength) that appear to limit the ‘destabilisation potential’ of experimentally trained memories in non-human animals (Milekic & Alberini, 2002; Suzuki et al., 2004; Vousden & Milton, 2017) do not necessarily preclude destabilisation of naturally acquired memories in humans, although they might constrain intervention effects. This provisional conclusion is promising for the development of such strategies as therapeutic interventions for threat-related and substance use disorders.

However, ESs varied widely across substance use and phobia/trauma studies. There was considerable variation in retrieval procedures (e.g. retrieval duration; timing of treatment relative to retrieval), the nature of reconsolidation interference strategies (i.e. the use of different drug classes and behavioural interventions), and participant characteristics. Specific combinations of these key variables likely influenced the efficacy of the overall reconsolidation modulating procedure and contributed to ES heterogeneity. As such, there continues to be uncertainty about optimal retrieval parameters and/or retrieval-dependent interference strategies required to interfere effectively with naturalistic maladaptive memories. Such memories are likely to constitute highly distributed traces and involve multiple memory systems (semantic, autobiographical-episodic, priming/implicit; varying in affective valence). This is clearly very different to the situation in studies of laboratory-trained memories, in which learning usually involves a limited set of stimuli from well-defined categories (e.g. sets of simple sensory stimuli as CSs) within a single context. Moreover, effective retrieval (reactivating) cues in laboratory studies are simply those that were used during training, whereas the nature of suitable retrieval cues for naturalistic memories is unclear. Given the uncertainty regarding suitable retrieval conditions for naturalistic memories, as well as the likelihood that such memories are more strongly entrained (over a long period) than experimentally acquired memories, it might be expected that ESs would differ between experimental and naturalist memories. It is therefore instructive to compare our ES estimates with those obtained in previous meta-analyses of reconsolidation studies of experimentally-acquired memories in humans. The results reported in two relevant meta-analyses on emotional (threat-related) memories are therefore considered (Kredlow et al., 2016; Lonergan, Olivera-Figueroa, Pitman, & Brunet, 2013) in relation to phobia/trauma studies.

### Phobia/trauma studies

Kredlow and colleagues (2016) examined changes in conditioned fear in studies employing the prototypical behavioural ‘retrieval-extinction’ procedure (Monfils et al., 2009). Their reported aggregate ES, relating to tests of ‘return of fear’ is not dissimilar to that for behavioural strategies (most commonly, retrieval-extinction) employed in the phobia/trauma studies reviewed here. Moreover, heterogeneity in the ESs in Kredlow et al (2016; Kredlow, personal communication) was similar to that of behavioural studies reviewed here (*I*^2^=52% and 43% respectively). By contrast, the aggregate ES reported here for phobia/trauma studies using pharmacological treatments (propranolol, mifepristone and sirolimus; *g*=0.71) was somewhat larger than that reported in a meta-analysis of studies of the effects of propranolol on memory for negatively valenced words and cue-induced fear responding (*g*=0.56; Lonergan, Olivera-Figueroa, et al., 2013). Both Lonergan et al., (2013) and the current analysis of pharmacological/phobia studies showed high levels of heterogeneity (*I*^2^=72%). These similarities underscores the potential for applying research findings from studies on experimentally acquired memories to naturalistic memories in clinical disorders. However, they also highlight the need for further research to identify common sources of heterogeneity in ESs in reconsolidation research.

As noted above, a single influential study (Soeter & Kindt, 2015a) contributed disproportionately to the moderate-large ES of pharmacological studies summarised here. Removal of this study drastically reduced the ES. However, given that this study produced the most pronounced, prolonged (1 year), and generalised (across behavioural and subjective-evaluative indices of fear memory) effects, it is worth considering the study features that might have been responsible for these particularly large and durable effects. It is noteworthy, for example, that Soeter & Kindt’s (2015a) study was the only one among the phobia/trauma studies to use propranolol as a pharmacological interference strategy (it was also the only pharmacological study on specific phobia rather than PTSD). This was based on their multiple previous demonstrations of reconsolidation impairing effects of propranolol on conditioned fear memories (Kindt, Soeter, & Vervliet, 2009; Sevenster et al., 2013; Sevenster, Beckers, & Kindt, 2012b; Sevenster et al., 2014; Soeter & Kindt, 2010, 2011, 2012a, 2012b, 2015a, but see Bos et al., 2014; Schroyens et al., 2017).

While these fear conditioning studies have demonstrated that relatively brief (<10 s) retrievals involving unreinforced presentation of a CS appear sufficient to reactivate conditioned fear memories, Soeter and Kindt (2015a) used a longer retrieval duration (2 min), as apparently required to reactivate chronologically remote memories (Suzuki et al., 2004). This was substantially longer than the retrievals used in the other (behavioural) reconsolidation-phobia studies (Bjorkstrand et al., 2016; Maples-keller et al., 2017; Shiban et al., 2015; Telch et al., 2017). Soeter & Kindt’s (2015a) retrieval procedure also involved a prediction error (participants expected that they would touch the spider during retrieval, but in fact, this did not occur). Finally, the retrieval procedure involved *in vivo* exposure to a spider (*cf.* Bjorkstrand et al., 2016; Maples-keller et al., 2017; Shiban et al., 2015), which, as a biological ‘prepared’ stimulus, might be considered to possess qualities of a US. Some researchers (e.g. Liu et al., 2014) have suggested that use of USs at retrieval results in a more generalised destabilisation (between all CSs and the US). This might explain the more generalised effects on fear responses in Soeter and Kindt (2015b). A close replication is now required to establish that this combination of factors (i.e. use of post-retrieval propranolol; medium duration retrieval and/or use of cues with US properties and/or incorporating a relevant prediction error at retrieval) reliably produces large and durable reconsolidation effects on naturalistic fear memories. Thereafter, studies might seek to determine if this combination is *required.*

The basic behavioural pharmacology of other neurotransmitter/neuromodulator systems is less well developed relative to the noradrenergic system. For example, despite the established role of glucocorticoid stress system in memory, disruption of which is implicated in psychological disorders (de Quervain, Schwabe, & Roozendaal, 2017), few basic behavioural studies have been conducted on its role in reconsolidation in humans. Moreover, endogenous and exogenous corticosteroids have a variety of distinct and opposing effects on memory (e.g. impairment of retrieval versus enhancement of (re)consolidation), depending upon, for example, timing of the glucocorticoid surge relative to retrieval, the number (or duration) of CS exposure at retrieval (e.g. Cai, 2006) and background levels of arousal (see Meir Drexler & Wolf, 2017). These multiple determinants of glucocorticoid effects might explain the conflicting results reported in existing human and non-human animal studies of glucocorticoids modulation of reconsolidation (de Quervain et al., 2017). It is unclear whether these considerations were relevant in the two Wood et al studies (2015 studies 2 and 3), both of which showed no statistical effect of mifepristone, a glucocorticoid receptor antagonist (although effects were in the predicted direction). In Wood et al (2015, Study 3), the authors additionally attempted to augment the impairing effects of mifepristone through pre-treatment with the NMDAR (glycine site) partial agonist, D-cycloserine. This strategy might be particularly relevant when long-term allostatic processes (Espejo et al., 2016) result in an enduring down-regulation of NMDAR (NR2B) subunits, which are required for memory destabilisation (Ben Mamou, Gamache, & Nader, 2006; Wang, De Oliveira Alvares, & Nader, 2009). As is evident from Figures 3 and 4b, DCS did not appear to affect mifepristone’s ability to interfere with reconsolidation of trauma memory.

The limited effects of mifepristone might be attributable to specific procedural factors in the two studies described in Wood et al (2015). For example, the use of individualised scripts likely introduced variability in retrieval duration across participants. In addition, apparently prolonged ‘script preparation’ procedures at retrieval (writing about two traumatic experiences from them same or different events and recalling subjective, visceral and muscular reactions associated with these experiences) might have engaged extinction rather than reconsolidation processes. These limitations in the extant research on drugs that downregulate glucocorticoid receptor activity do not allow firm conclusions to be drawn about their application as reconsolidation-interfering treatments.

Unlike the role of the noradrenergic and glucocorticoid systems, little is known about the effects of manipulating the mTOR pathway on any aspect of memory functioning in humans. While rapamycin blocks fear-related memory reconsolidation in rodents (e.g. Blundell, Kouser, & Powell, 2008) this capacity has not yet been established in experimental studies of emotional learning and memory in humans. This makes it difficult to interpret the limited efficacy of rapamycin reported in (Surís et al., 2013). In addition to the long retrieval duration (up to 75 min) used in that study, the pharmacokinetic profile of rapamycin (i.e. its central bioavailability after a single 15 mg oral dose) relative to the timing of reactivation (assuming this actually occurred) may not have been optimal.

In contrast to the ES estimate from pharmacological studies of phobia/trauma, the population ES from studies of post-retrieval behavioural strategies was small (less than half that obtained from pharmacological studies) and was not reliably different from 0. Despite four of the five ESs favouring the Retrieval + Treatment group, the effect was skewed towards a smaller value by the Shiban et al (2015) study. The ES in that study was based on a spontaneous recovery test performed one day after retrieval-extinction. If there is a ‘sleeper effect’ on BAT performance, as found for declarative aspects of fear by Soeter and Kindt (2015a; note these authors tested behavioural approach for the first time 11 days after treatment with propranolol), retrieval-dependent effects might not have been evident after such a short interval. However, in contrast to Soeter and Kindt (2015a), Shiban et al. (2015) reported no evidence for enhanced effects of retrieval-extinction on declarative fear after a long follow-up period (6 month). In addition to differences in the retrieval procedures between these two studies, (non-significantly) higher baseline levels of fear, heart rate and skin conductance, as well as lower baseline approach behaviour in the retrieval-extinction group (relative to the control group in Shiban et al., 2015), might have contributed to a limited effect of extinction in the former group.

One notable finding among the behavioural phobia/trauma studies was the relatively immediate reduction in fear responding (expectancy and peak fear) in the Retrieval + Treatment (extinction) group relative to a Treatment + Retrieval control group in Telch et al. (2017). These authors consider a number of explanations for this unexpected early effect (e.g. the occurrence of prediction error or increased noradrenergic activity resulting from the retrieval trial). However, it should also be noted that the interval between retrieval and the first extinction trial was especially long in this study (30 min). While the use of a delay between retrieval and the interference strategy is a common feature of behavioural reconsolidation studies (although the interval is usually only 10 min), the rationale for employing such a delay is unclear and may simply be a carryover from the procedure used in the first studies of retrieval-extinction in rodents and humans (Monfils et al., 2009; Schiller et al., 2009). Indeed, without employing *high cognitive load tasks* during the retrieval-interference interval (e.g. Das et al., 2015b; Das et al., 2018; Hon et al., 2015), there is the potential for ongoing cognitive engagement/rehearsal following exposure to the reminder cue, possibly initiating extinction. Not only would this be expected to limit longer-term reconsolidation-dependent effects, but also suggests an alternative explanation for the apparent retrieval dependent enhancement of early extinction reported by Telch et al (2017).

### Substance use studies

Despite the small number of studies of laboratory-based reward-memory reconsolidation in humans (e.g. Xue et al., 2017; Zhao et al., 2011), there is substantial evidence for reconsolidation modulation in rodent models of maladaptive reward/addiction. Meta-analytic findings on appetitive-reward memory in non-human animals suggest that the ES associated with reconsolidation interference using NMDAR antagonism is substantially larger than that for β-adrenergic antagonists (Das et al., 2013). This might explain the relatively modest and short-lived effects of propranolol reported by Saladin et al. (2013). Alternatively, the retrieval procedure in (Saladin et al., 2013) was relatively protracted (2 × 10 min of *in vivo* and video cue exposure to cocaine cues), which might have limited plasticity through activation of extinction or generating a limbo state. The other study that used propranolol (Pachas et al., 2015) showed a *negative* ES (the propranolol group showing *higher* levels of craving relative to Retrieval + No Treatment). It should be noted however, that the latter study used a script preparation protocol inspired by the same studies that informed the Wood et al (2015) retrieval procedures. Again, while the duration of the retrieval procedure was not stated in Pachas et al (2015), it is likely that it was also prolonged (and likely to vary between participants). As such the potential for extinction /limbo state processes is again relevant. Since craving was *higher* in the Retrieval + Treatment group, it is possible that extinction consolidation was *impaired* by propranolol relative to the placebo group (Cahill, Pham, & Setlow, 2000).

Excluding Pachas et al (2015), the remaining two pharmacological studies (Das et al., 2015a; Saladin et al., 2013) showed no evidence of a combined effect consistent with reconsolidation interference. Based on the larger effects of NMDAR-versus β-adrenergic antagonists on reward memory reconsolidation (Das et al., 2013), Das et al. (2015a) examined the NMDAR antagonists, memantine, but found no evidence for an effect consistent with reconsolidation blockade. It is unclear whether the typical therapeutic dose (10 mg) and route of administration (oral) of memantine is suitable for blocking reconsolidation in humans. Indeed, memantine has slow absorption kinetics, and relatively low selectivity for the most abundant central NR2A NMDAR subunit (Ogden & Traynelis, 2011), which are involved in restabilisation (Milton et al, 2013) giving rise to uncertainty about whether suitable reductions in NMDAR activity were achieved in the post-retrieval period.

In contrast to pharmacological strategies, behavioural methods for interfering with reconsolidation showed more promise in the case of substance use. Indeed the ES associated with behavioural studies appeared to be significantly larger than that of pharmacological studies, although this statistical finding needs to be treated with caution given the small number of studies and lack of precision in the population ES estimate of pharmacological studies. Of the four behavioural-substance use studies, two employed post-retrieval cue exposure (‘retrieval-extinction’; Germeroth et al., 2017; Xue et al., 2012, one counterconditioning Das et al., 2015b, and one, cognitive reappraisal; Hon et al., 2015). All showed positive ESs on measures of craving, and the overall ES was moderate-large, with minimal heterogeneity. It is noteworthy that while the nature of the reactivating cues varied between studies (e.g. drug use video, in vivo cues, drug pictures or a combination of these) all of the behavioural-substance use studies used the same retrieval duration (5 min), along with a 10 min interval between termination of retrieval and start of the behavioural strategy.

In addition, it is noteworthy that all of the substance use behavioural-interference studies reported significant effects on more than one outcome. Indeed, Das et al. (2015b), reported results consistent with a comprehensive rewriting of affective, attentional and cognitive aspects of alcohol related memories in heavy drinkers. However, the follow-up period for this study – along with Hon et al. (2015) was relatively short (1 week), whereas the other two studies tested participants at 1 month (Germeroth et al., 2017) and 6 months (Xue et al., 2012). Three of the studies examined changes in substance use behaviour (Das et al., 2015b; Germeroth et al., 2017; Hon et al., 2015), but only one of these showed significant changes in drug (cigarette) use (Germeroth et al., 2017). The latter study, along with the (Xue et al., 2012) study used a two session treatment protocol (two Retrieval + Treatment session). As such, there were considerable differences across studies in terms of longest follow-up time-point and/or effects on behaviour. It remains to be determined whether counterconditioning and cognitive reappraisal result in sustained effects on craving (and attentional bias and effective responding to alcohol: Das, et al., 2015b; and semantic memory for alcohol, Hon et al., 2015, and whether behavioural effects might also emerge after a longer delay.

#### Limitations

The effects reported here are based on relatively small numbers of studies in each category of disorder (these were further reduced in the moderator analyses). In addition, most studies reviewed here had small sample sizes. Moderation was only examined for a small number of covariates in the current analysis. However, assuming detailed methodological reporting in future studies, meta-analysis/regression based on a larger number of studies with greater variability in retrieval variables might prove to be a particularly effective way of establishing the role of retrieval parameters (e.g. retrieval duration; use of prediction error at retrieval) in successful memory reactivation. The alternative-parametric variation of these retrieval parameters in experimental studies-would require unrealistically large sample sizes due to the number of potential factors and levels of these key variables.

In contrast to the array of outcomes reported in the reviewed studies, our analysis focused on a narrow set of pre-determined outcomes (primarily trauma symptoms in studies of trauma-exposed individuals, behavioural approach or subjective fear in phobia studies and craving in the case of substance use studies). We opted to base our ES calculations on these outcomes rather than those preferred by authors/reported as statistically significant in publications. However, it is possible that quite different results would be achieved if only the significant results reported by study authors (either singly or as composites of multiple significant outcomes) were used to determine ESs. This is not necessarily a limitation, as our intention was to reduce the potential for bias in cases where multiple outcomes were reported but none were pre-specified as primary.

### Recommendations for future research

A strength of the reviewed studies was their tendency to recruit participants in line with the relative gender prevalence of disorders in question. However, it should be noted that recent evidence suggests that men and women may be differentially susceptible to some reconsolidation interfering treatments. In particular, Drexler and colleagues (2016) showed that whereas men showed retrieval-dependent weakening following hydrocortisone, this effect was absent in women. It is currently unclear whether this finding is specific to glucocorticoid modulation of reconsolidation, rather than reflecting a general insensitivity to reconsolidation interference in women. Indeed, the latter seems highly unlikely given the very large effects seen with propranolol seen in Soeter and Kindt (2015a), whose sample consisted almost exclusively of women (91%). No individual study that we are aware of has yet examined gender moderation in (sub)clinical populations, although this seems a particularly important factor to consider if reconsolidation-interference is to be used clinically. It should be noted provisionally, that our moderation analysis did not suggest an effect of gender.

Among the pharmacological studies, there was no common use of a single reconsolidation-interfering drug. Although propranolol was most commonly studied, this amounted to only two studies in the substance use category and one study in the phobia/trauma grouping. As such, it remains unclear whether one drug class category might be more effective in preventing restabilisation than others. Despite strong evidence from studies with non-human animals, only one of the reviewed studies examined an NMDAR antagonist. Given the central role of glutamatergic neurotransmission in learning and memory, further research on the effects of NMDAR antagonist effects on reconsolidation in humans seems to be a special priority. On the other hand, clinical studies should be preceded by more basic psychopharmacological studies in order to determine the importance of drug timing (relative to retrieval) and route of administration. This is particularly important given the potential for iatrogenic effects of NMDAR antagonists (i.e. paradoxical strengthening of maladaptive memories in some contexts; Honsberger, Taylor, & Corlett, 2015).

Despite its reported importance of memory destabilization, few of the reviewed studies examined the role of prediction error (*cf*. Das et al., 2015b; Das et al., 2015a; Hon et al., 2015; Soeter & Kindt, 2015a). As noted previously, if there is a requirement for an optimal learning signal at retrieval, those studies showing beneficial effects of Retrieval + Treatment in the absence of prediction error, might in fact represent the lower bound of efficacy that could be achieved during reconsolidation modulation. As such, tailoring retrieval procedures to maximise PE may bolster the likelihood that reconsolidation can be leveraged for clinical benefit.

Overall, our findings suggest that reconsolidation-interference is worth pursuing as a clinical strategy. However, the multiple sources of uncertainty regarding determinants of efficacy suggest further basic research is needed to ensure that studies with clinical populations are as informative as possible.

